# Switching toxic protein function in life cells

**DOI:** 10.1101/430439

**Authors:** Frederik Faden, Stefan Mielke, Nico Dissmeyer

**Affiliations:** Independent Junior Research Group on Protein Recognition and Degradation, Leibniz Institute of Plant Biochemistry (IPB), Weinberg 3, D-06120 Halle (Saale), Germany; ScienceCampus Halle – Plant-based Bioeconomy, Betty-Heimann-Str. 3, D-06120 Halle (Saale), Germany; CSL Behring AG, Wankdorfstrasse 10, CH-3014 Bern, Switzerland; Jasmonate Signalling Group, Leibniz Institute of Plant Biochemistry (IPB), Weinberg 3, D-06120 Halle (Saale), Germany

**Keywords:** cell ablation, cell death, conditional alleles, molecular farming, N-end rule, plant-made pharmaceuticals/biopharming, protein degradation, trichomes

## Abstract

Toxic proteins are prime targets for molecular farming and efficient tools for targeted cell ablation in genetics, developmental biology, and biotechnology. Achieving conditional activity of cytotoxins and their maintenance in form of stably transformed transgenes is challenging. We demonstrate here a switchable version of the highly cytotoxic bacterial ribonuclease barnase by using efficient temperature-dependent control of protein accumulation in living multicellular organisms. By tuning the levels of the protein, we were able to control the fate of a plant organ *in vivo*. The on-demand-formation of specialized epidermal cells (trichomes) through manipulating stabilization versus destabilization of barnase is a proof-of-concept for a robust and powerful tool for conditional switchable cell arrest. We present this tool both as a potential novel strategy for the manufacture and accumulation of cytotoxic proteins and toxic high-value products in plants or for conditional genetic cell ablation.

## INTRODUCTION

The low-temperature degron cassette (lt-degron) fused to a protein of interest (POI) can efficiently control its accumulation or degradation in a temperature-dependent manner. The technique is based on altering the *in vivo* stability of the lt-degron:POI fusion protein by using a temperature stimulus and relies on the proteostatic machinery of the N-end rule pathway of protein degradation (Dissmeyer et al., 2018) for fast removal from the cell. The lt-degron:POI fusion can thus be efficiently accumulated at a permissive temperature or removed from the cell at a restrictive temperature generating an artificial temperature-sensitive variant of the POI (Faden et al., 2016; Dissmeyer, 2017). Opposed to many other systems for conditional protein expression and gene regulation, the lt-degron approach relies on posttranslational oinzerference by protein degradation of the entire fusion protein and is applicable in living multicellular organisms, in both plants and animals. By this, it offers tight control over the levels of active/functional protein which directly correlates to the temperature stimulus used to induce the system.

Molecular farming, the generation of pharmacologically active or biotechnologically usable compounds in plants, is an emerging topic especially for the production of peptides and proteins requiring special glycosylation patterns impossible to generate in classical fermenter-based expression systems using microorganisms like bacteria (Stoger et al., 2014). Various cytotoxic peptides are under investigation as highly potent cancer therapeutics (Boohaker et al., 2012). Some toxic proteins, such as the mistletoe lectins possess anti-cancer properties when administered orally (Pryme et al., 2007) and recombinantly produced immunotoxins are in clinical and pre-clinical evaluation for cancer treatments (Kelly et al., 2012; Madhumathi and Verma, 2012). So far, lectin proteins from plants as well as from animals have mainly been efficiently produced in *E. coli* with moderate yields (Lam and Ng, 2011), however, the production of toxic peptides and proteins in plants is still challenging also due to difficult transgene maintenance. So far, production of phytotoxic peptides, such as e.g. antimicrobial peptides, has been difficult and needed either further modification of the peptide (Company et al., 2014) or was achieved through vigorous control of transgene expression through e.g. inducible promoters (Company et al., 2014). The lt-degron technique offers tremendous advantages over inducible promoters due to easier control and non-aversive effects of the stimulus of elevated ambient temperature versus chilling.

The bacterial ribonuclease barnase (BAR), a potent, non-specific RNAse secreted by the soil bacterium *Bacillus amyloliquefaciens* (Buckle and Fersht, 1994) possesses anti-cancer properties (Edelweiss et al., 2008) and has also been used as a potent tool for plant breeders to create male sterile mutants in tobacco, *Arabidopsis*, and wheat (Glimcher and Sparks, 1992; Burgess et al., 2002; Gils et al., 2008). Additionally, it has been exploited as a cell-ablation tool to destroy inflorescences to prevent spread of transgenes from birch, tobacco, and *Arabidopsis* (Lannanpaa et al., 2005). Other applications include a system to defend potato plants against the pathogen *Phytophthora infestans* (Strittmatter et al., 1995) and a conditional cell ablation tool in mammalian cell culture (Leuchtenberger et al., 2001). Due to the conserved mechanism of barnase activity as a non-specific RNase, it is considered to be highly cytotoxic in literally all organisms, highlighting the broad spectrum of possible applications for a controllable ribonuclease.

Cell ablation has been proven to be a powerful tool to study the effects of elimination of entire cell populations in a biological context. Conditional as well as constitutive techniques have been extensively used in the animal field, e.g. to map cell populations in the murine nervous system where the natural resistance of some mouse cell types against the highly cytotoxic *Diphtheria* toxin chain A (DT-A) has been exploited. By controlling the expression of a DT-A receptor, cells were indirectly but efficiently ablated upon exogenous administering of DT-A (Saito et al., 2001; Hermand, 2006). DT-A was also used extensively to induce cell death in yeast, fruit flies, and plants (Bellen et al., 1992; Guerineau et al., 2003). Another example is a regeneration study in zebra fish through the guided, enzymatic conversation of a prodrug (precursor drug) into an active, DNA-toxic compound by a bacterial nitroreductase (Curado et al., 2007). Another system, albeit not applicable in plants, is the auxin-controlled ‘suicide module’ based on rapid degradation of the inhibitory chaperone ICAD. Its degradation enabled caspase activity inducing cell death (Samejima et al., 2014).

Cell ablation by toxic proteins needs to be tightly controlled in a conditional rather than constitutive manner. Otherwise, the host organism or expressing tissue will be irreversibly destroyed. However, most conditional, non-constitutive, cell ablation protocols rely on the addition of exogenous compounds such as hormones or other small molecules triggering cell death. Depending on the experimental conditions, these systems struggle with uneven induction of the system, low penetration or metabolic secondary effects of the inducing agents. Moreover, the systems have intrinsic delays in response due to the nature of induction of transcription followed by translation of target protein. The protein accumulation is irreversible which makes control at the level of acute toxicity namely the protein level, impossible.

Here we report a strategy to control barnase protein levels via temperature and therefore turn on/off its activity. This allows generating a *“phenotype on demand"* and results in conditional organ formation versus ablation in intact, living plants. We combined the lt-degron with the cytotoxic barnase protein and expressed the resulting lt-BAR construct in leaf hairs (trichomes), which are specialized single cells of the epidermis, under control of the trichome-specific *TRIPTYCHON* (*TRY*) promoter (Pesch and Hulskamp, 2011).

This setup allowed controlling trichome cellular fate in a temperature-dependent manner and serves as a proof-of-concept for conditional cell ablation. This new tool also paves the way towards a more efficient molecular pharming of cytotoxic proteins *in planta*.

## MATERIALS AND METHODS

### Cloning and DNA work

DNA cloning was performed following standard procedures using *Escherichia coli* strain DH5a (Invitrogen). The constructs were generated by fusion PCR (*Pfu* poly-merase, fermentas) using the following primer combinations. DNA amplicons were purified with ExoSap-IT (USB) preceding fusion PCRs. All fusions were flanked by Gateway *attB1/attB2* sites and recombined by BP reactions into *pDONR201* (Invitro-gen).

The lt-degron cassette (K2) was amplified from *pEN-L1-K2-L2* (Addgene ID: 80684)^1^ using primers TMV_att_frw (GGGGACAAGTTTGTACAAAAAA-

GCAGGCTTActcgagctgcagaattactatttacaattac) and K2barn_f1_rev (tgtgccatagcac-cagcaccagcgtaa).

The lt-degron K2 starts with a 3’ tobacco Omega (Ω) leader sequence for translation enhancement from *pRTUB8*, a derivative of *pRTUB1* (Bachmair et al., 1990; Bachmair et al., 1993). The leader contains the 20 nucleotides upstream of the start codon of tobacco mosaic virus strain U1(Gallie et al., 1987) and is followed by a co-don-optimized synthetic human *Ubiquitin* gene(Ecker et al., 1987), a triplet for the destabilizing bulky and hydrophobic amino acid Phe (F), a short linker peptide (translates to HGSGI)(Bachmair and Varshavsky, 1989) to further expose the N-terminal residue, a temperature-sensitive mouse dihydrofolate reductase sequences (*DHFR^ts^*) triggering the protein unfolding response, a triple hemagglutinin (HA)-tag (HAT) for immunodetection. Phe is used as destabilizing N-terminal residue, because it was shown to be recognized by the *Arabidopsis* N-end rule pathway(Bachmair et al., 1993; Potuschak et al., 1998; Stary et al., 2003; Faden et al., 2016; Mot et al., 2018). HAT was amplified from *pSKTag3SUM6* (kind gift of Andreas Bachmair, Max F. Perutz Laboratories, Vienna, Austria). The DHFR moiety of K2 is derived from *pJH10^mut^* (*pJH23*), containing a mutated mouse *DHFR^T39A,E173D^* (Gowda et al., 2013). An Entry clone containing *K2:TTG1* was used as a template for *K2* (Faden et al., 2016). BARNASE, including the artificial intron, was amplified from *pICH43601* (kind gift of Sylvestre Marillonnet, IPB Halle, Germany) using primers K2barn_f2_frw (ctggtgc-tatggcacaggttatcaacacgtt) and K2barn_f2_rev (GGGGACCACTTTGTACAAGAAA-GCTGGGTAttatctgatttttgta). Primers K2barn_f1_rev and K2barn_f2_frw contained each an eight basepair overhang to the respective other fragment. A fusion PCR was carried out using primers TMV_att_frw and K2barn_f2_rev. The fusion product was flanked with BP compatible Gateway recombination sites, subcloned into *pDONR201* (Invitrogen) and the *pENTR::lt-BAR* Entry clones sequenced after a BP reaction. For stable expression in transgenic plants, *pENTR::lt-BAR* was recombined in an LR reaction into the *attR* site-containing binary Gateway destination vector *pAM-PAT:ProTRY:GW* (Addgene ID: 79755 (Pesch and Hulskamp, 2011)) yielding *pAM-PAT:ProTRY::lt-BAR*. These Gateway-compatible Destination vectors are derived from *pAM-PAT-MCsS* (multiple cloning site; GenBank accession number AY436765), derivative of *pPAM* (GenBank accession number AY027531) (Rademacher et al., 2002). These backbones carry the *Streptomyces hygroscopicus bar* gene that translates to phosphinothricin-N-acetyltransferase (PAT) as plant selection marker conferring resistance against PPT. This vector was used for Agrobacterium-mediated transformation of Col-0 wildtype plants.

### Plant work and growth conditions

*Arabidopsis thaliana* (L.) Heynh. plants were grown either on a steamed (sterilized for 3h at 90°C) soil mixture (Einheitserde Classic Kokos (45% (w/w) white peat, 20% (w/w) clay, 15% (w/w) block peat, 20% (w/w) coco fibers; 10–00800–40, Einheitserdewerke Patzer); 25% (w/w) Vermiculite (grain size 2–3 mm; 29.060220, Gärtnereibedarf Kamlott); 300–400 g of Exemptor (100 g/kg thiacloprid, 802288, Hermann Meyer)) per m^3^ soil mixture or aseptically *in vitro* on plastic Petri dishes containing ½ Murashige & Skoog (MS) (2.16 g/l MS salts, Duchefa Biochemicals, M0221; 0.5% Glu sucrose; 8 g/l phytoagar, Duchefa Biochemicals, P10031; pH to 5.6 - 5.8 using KOH). Aseptic culture was done or under long-day regime (16/8 hours light/dark) in growth cabinets.

Seeds for soild-grown plants were germinated after stratification of four to five days at 4°C in the dark and plants grown under standard long-day (16/8 hours light/dark) or short day (8/16 hours light/dark) greenhouse conditions between 18 and 25°C. For strictly controlled development such as during temperature shift experiments, plants were grown either in growth cabinets (AR-66L2 and AR-66L3, Percival Scientific, CLF PlantClimatics) or walk-in phyto chambers (Johnson Controls, equipped with ESC 300 software interface) at a humidity of 60% depending on the requirements and watered with prewarmed or precooled tap water.

Transgenic, Basta resistant plants were selected *in solium* in cotyledon stage by spraying 150 mL per tray of a 1:1000 dilution of Basta (contains 200 g/L glufosinate-ammonium; Bayer CropScience) in tap water which was repeated three times in a two-days interval.

For aseptic culture, dry seeds were sterilized with chlorine dioxide gas produced from 75% *Eau de Javel* (FLOREAL Haagen) and 25% HCl. Selective MS media contained 10 mg/L DL-phosphinotricin (PPT, Basta, glufosinate-ammonium, sc-235254, Santa Cruz Biotechnology), 50 mg/L kanamycin sulfate, 5.25 mg/L sulfadiazine sodium salt (Sigma-Aldrich, S6387), or 20 mg/L hygromycine B (Duchefa Biochemicals, H0192). Plants used in this study were all in the background of the Columbia-0 accession (Col-0) and either wild-type plants or an ethyl methanesulfonate (EMS) mutant for *PRT1* (*prt1* HygS or *prt1–1*, kind gift of Andreas Bachmair, Max F. Perutz Laboratories, Vienna, Austria),(Bachmair et al., 1993; Potuschak et al., 1998; Stary et al., 2003)

For controlled environment experiments plants were cultivated in growth cabinets (AR-66L2 or AR-66L3, Percival Scientific, CLF PlantClimatics). For experiments needing less stringent environmental conditions plants were cultivated in the greenhouse.

### Stable plant transformation and selection of transformants

The binary plant Expression vectors were retransformed into *Agrobacterium tumefaciens GV3101-pMP90RK* (*C58C1 Rif^r^ Gm^r^* *Km^r^*)(Koncz and Schell, 1986) to obtain a bacterial transformation suspension. The identity of the *Agrobacterium* strains was verified by backtransformation of isolated plasmid into *E. coli* DH5α and at least three independent analytical digestions. All constructs were transformed by a modified version of the floral dip method (Dissmeyer and Schnittger, 2011).

Individual T1 (generation 1 after transformation) transgenic plant lines were pre-selected with Basta or kanamycin as described above. To exclude lines showing position effects, e.g. by disrupting essential genes by the construct T-DNA, the number of insertion loci was determined in a segregation analysis in the T2 generation and only transgenic plants carrying one single insertion locus were further used. Standard lines were established by isolating T3 plants homozygous for the transgene. Independent representative reference lines displaying a typical conditional phenotype upon temperature up- and downshifts were used in the final experiments. Standard lines were established by isolating T3 plants homozygous for the transgene. In order to identify responsive transgenic lines, we prescreened by developmental and histological phenotype.

### Transient transfection of tobacco

For transient transformation of ***Nicotiana benthamiana* (tobacco)**, leaves of five-week-old plants were infiltrated with *Agrobacteria GV3101-pMP90RK*, carrying binary plant expression vectors. *Agrobacteria* were grown to the stationary phase overnight in 10 mL of YEB medium and pelleted at 5000 x g at RT. The pellet was washed once in 10 mL of infiltration buffer (10 mM MES pH 5.6, 10 mM MgSO_4_, 100 μΜ ace-tosyringone (D134406, Sigma)) and subsequently resuspended in 10 mL of the same buffer. *Agrobacteria* strains were always co-infiltrated together with the *Agrobacterium* strain GV3101 expressing *p19* of tomato bushy stunt virus from the vector *pBIN6Ip19* to suppress post-transcriptional gene silencing and increase ectopic gene expression (Voinnet et al., 2003). Prior to infiltration, bacteria suspensions were adjusted to an OD_600_ of 0.5. Bacteria suspensions containing the lt-BAR transgene were mixed with *p19* bacteria suspension in a 1:1 ratio (final OD_600_ = 0.5) and used for transformation. Bacteria suspensions were then infiltrated into the epidermis on the lower side of the tobacco leaf. Infiltrated areas were marked on the upper side of the leaf with a permanent marker. For an easier infiltration procedure, plants were watered and transferred to standard greenhouse LD conditions the day before in order to allow plant stomata to open. To allow efficient transformation and expression plants were kept for 48 h in the greenhouse, before applying temperature shift experiments by transferring them into a growth cabinet at either permissive or restrictive growth conditions. For one data point, 15 - 20 leaf discs of 5 mm diameter were harvested from infiltrated areas and snap frozen in liquid nitrogen. Extraction was performed using RIPA buffer as mentioned below.

### Microscopy and documentation of plants

Photographs of *in vitro* cultures and histological stainings were taken with a stereo microscope (Stemi 2000-C) equipped with a Zeiss CL 6000 LED illumination unit, and a video adapter 60 C including an AxioCam ERc 5s digital camera (all from Carl Zeiss MicroImaging). Trichome visualization through polarized light microscopy was carried out as described previously (Gudesblat et al., 2012; Pomeranz et al., 2013) with the only change that destaining was achieved with a saturated chloral hydrate solution (dissolve 1 kg in 400 ml of H2O). Trichomes were visualized using a Nikon AZ100 zoom microscope equipped with a Nikon DS-Ri2 color camera. Microscopy images of trichomes and DAPI stained nuclei were obtained using the AxioImager system (Zeiss) equipped with two cameras (AxioCam MRm und AxioCam MRc5).

### Protein extraction and western blot analysis

Plant tissue (*Arabidopsis:* leafs or seedlings, tobacco: five leaf discs) was collected in a standard 2 mL reaction tube containing three Nirosta stainless steel beads (3.175 mm; 75306, Mühlmeier), snap frozen in liquid N_2_ and stored at −80°C. Material was ground frozen using a bead mill (Retsch, MM400; 45 s, 30 Hz) in collection microtube blocks (adaptor set from TissueLyser II, 69984, Qiagen; (Faden et al., 2016; Dissmeyer, 2017). Tissue was lysed using radioimmunoprecipitation assay (RIPA) buffer (50 mM Tris-Cl pH 8, 120 mM NaCl, 20 mM NaF, 1 mM EDTA, 6 mM EGTA, 1 mM benzamidine hydrochloride, 15 mM Na_4_P_2_O_7_, and 1% Nonidet P-40 supplemented with freshly added EDTA-free Complete Protease Inhibitor cocktail, Roche). The protein content of the samples was determined using a DirectDetect infrared spectrophotometer (MerckMillipore). Equal protein amounts were resolved by 12% SDS-PAGE. Blotting was done onto polyvinylidene fluoride (PVDF) transfer membrane by semi-dry blot using a Trans-Blot SD semi-dry electrophoretic transfer cell (170–3940, Bio-Rad).

Equal loading and general protein abundance was confirmed by staining of the blotted and probed membranes after immunostaining with Coomassie Brilliant Blue G250 (CBB G150). Immunodetection was done using mouse monoclonal anti-HA epitope tag antibody (HA.11, clone 16B12: MMS-101R, Covance, 1:1000) and HRP conjugated secondary antibodies which were detected by enhanced chemiluminescence (ECL) using ECL SuperSignal West Pico or Femto (34087 or 34096, Pierce) followed by exposure on autoradiography film. Western data were confirmed by analysis of at least three biological replicates.

### Transcript analysis

Two-week-old seedlings were collected and snap-frozen. After milling, about 50 mg of tissue was used for RNA extraction with RNeasy Plant Mini Kit (Qiagen). RNA was measured with a spectrophotometer and quality assessed via agarose gel electrophoresis. For first-strand cDNA synthesis, 500 ng of total RNA were used with an equimolar mixture of four oligo(dT) primers (CDSIII-NotIA/C/G/T (Faden et al., 2016)) and RevertAid H Minus Reverse Transcriptase (Thermo Scientific). 1 μL of cDNA was used for PCR analysis using self-made *Taq* in two reactions per sample; one with generic degron-specific primers (DHFR_frw/DHFR_rev) to test transcript levels of the transgene and one with intron-spanning primers EF1ss/EF1as for *ELONGATION-FACTOR1* (*EF1*) as a housekeeping gene (amplicon sizes: genomic 810 bp; cDNA 709 bp). Both PCRs were run for 30 cycles.

## RESULTS

### Conditional toxicity of ribonuclease leads to specific cell ablation in stable transgenic lines

The lt-degron cassette was fused to the coding sequence of barnase (BAR) that contains an artificial intron efficiently suppressing expression of functional toxic protein in bacteria due to possible promoter leakiness (Gils et al., 2008).

When expressed in trichomes of wildtype plants under the control of the *TRY* promoter (*ProTRY*), lt-BAR resulted in efficient ablation of trichomes. When plants were grown at the permissive temperature of 14°C, leaves appeared completely glabrous, as known from plants with mutated transcriptional regulators needed for trichome formation such as e.g. TRANSPARENT TESTA GLABRA1, GLABRA1 and 3, or CAPRICE (Hulskamp, 2004). Constitutive growth under the restrictive temperature of 28°C led to a destabilization of lt-BAR and allowed trichomes to form on newly emerging leaves (**Figure 1a**). Next, we evaluated the efficiency of the organ ablation through analysis of trichomes within the leaf epidermis. We used polarized light microscopy to analyze trichome spacing and number on true leafs. The wild type developed trichomes under both permissive and restrictive conditions whereas no trichomes could be spotted on mature leaves of *ProTRY::lt-BAR* plants when grown at the permissive temperature (**Figure 1b**). In contrast, these *lt-BAR* expressing plants fully retained trichomes when grown under restrictive conditions. This demonstrates the high efficiency and temperature-sensitivity of the function of the lt-BAR fusion protein as well as its effect on cell fate and organ formation. At the restrictive temperature, leaves of *ProTRY::lt-BAR* expressing plants were indistinguishable from wild type plants demonstrating an efficient degradation of the lt-BAR fusion protein to an extend where the barnase protein is unable to elicit a phenotype (**Figure 1b**).

**Figure 1.**
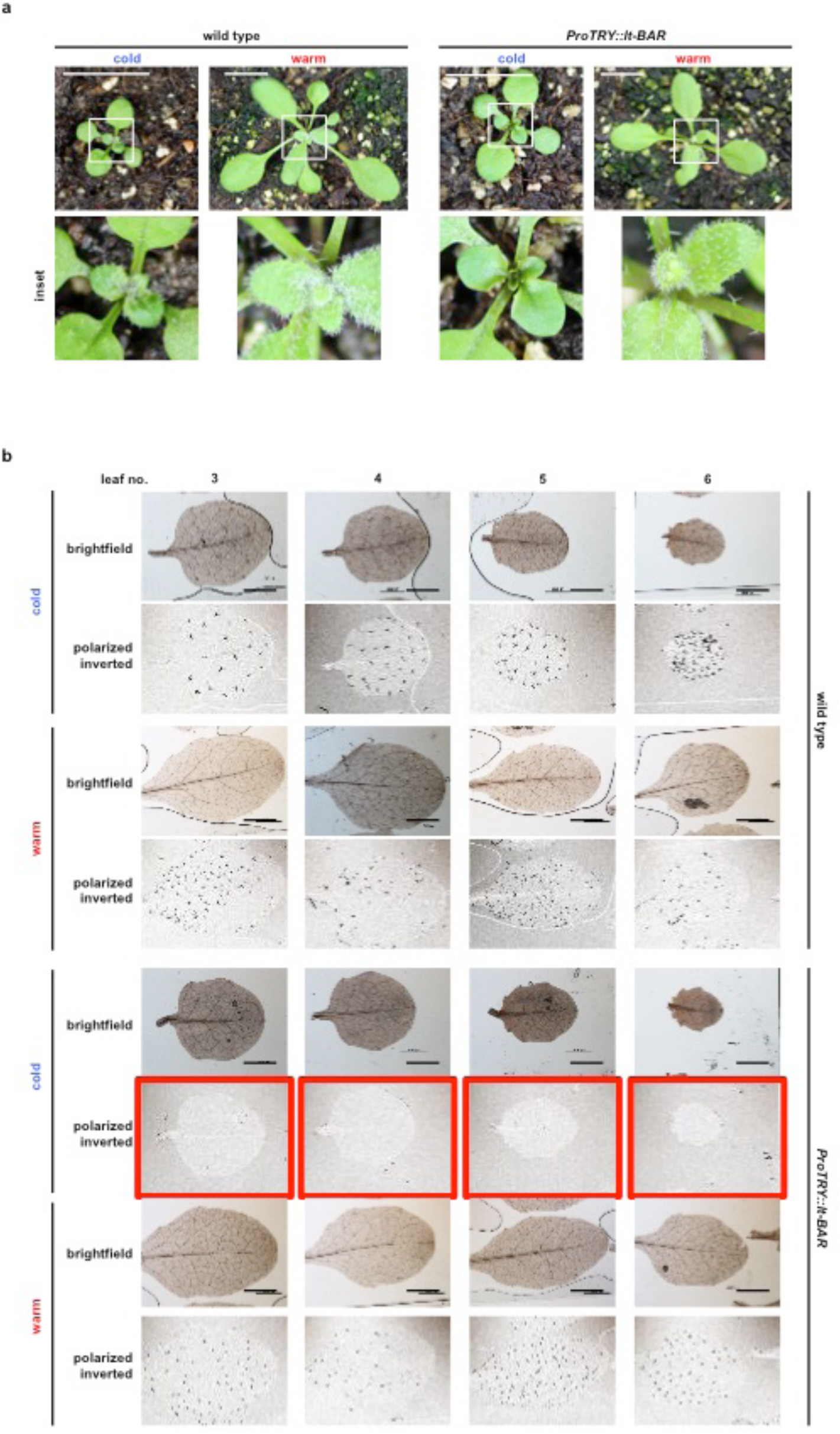
Conditional function of lt-BAR mediates robust temperature-dependent ablation of specific cells. **(a)** Plants expressing *ProTRY::lt-BAR* and the wild type grown for two weeks at permissive (cold) and restrictive (warm) temperatures. Leaves of transgenic plants appear glabrous at the permissive temperature. Size bar, 1 cm. **(b)** In-depth analysis of true leaves of *ProTRY::lt-BAR* and wildtype plants using polarized light microscopy. Conditionally cold or warm grown plants three weeks after germination. Cold grown transgenic plants show glabrous leafs (red boxes). Size bar, 500 μm.

### Transient expression of lt-barnase in tobacco leaves

The lt-degron, as a conditional protein ‘shut-off’ technique, relies on a temperature-sensitive switch between POI depletion and accumulation on protein level without altering transcript levels (Faden et al., 2016; Dissmeyer, 2017). Therefore, we tested conditional accumulation of lt-BAR upon shift of *ProTRY::lt-BAR* expressing *Arabidopsis* plants to permissive temperature. Although the cell-ablation phenotype was visible (**Figure 1a,b**), detection of lt-BAR protein with standard western blotting techniques gave only variable and unsatisfactory results. Because also detection of *lt-BAR* transcript of *ProTRY::lt-BAR* by semi-quantitative RT-PCR proved to be difficult, which is possibly due to very low levels of transcript as an effect of the additional RNAse activity from barnase, we transiently expressed the lt-BAR construct under the control of the *UBIQUITIN10* promoter (*ProUBQ10*) in *Nicotiana benthamiana* (tobacco) leaves. Surprisingly, the expression led to an almost temperature-independent phenotype with developing necrosis and chlorosis of the leaves at both permissive and restrictive temperature. The phenotype even appeared to be stronger at the restrictive temperature. This might be explained through an override of the degradation system in regard to lt-BAR-dependent degradation, most likely being a result of expression from the strong *ProUBQ10* combined with a possibly higher catalytic activity of the lt-BAR protein at the higher temperature (**Figure 2a**). We tried once more to demonstrate the connection between the phenotype and the abundance of lt-BAR fusion protein by western blotting and semi-quantitative RT-PCR and could obtain transcript data clearly linking the observed phenotype to the expression of lt-BAR (**Figure 2b**).

**Figure 2.**
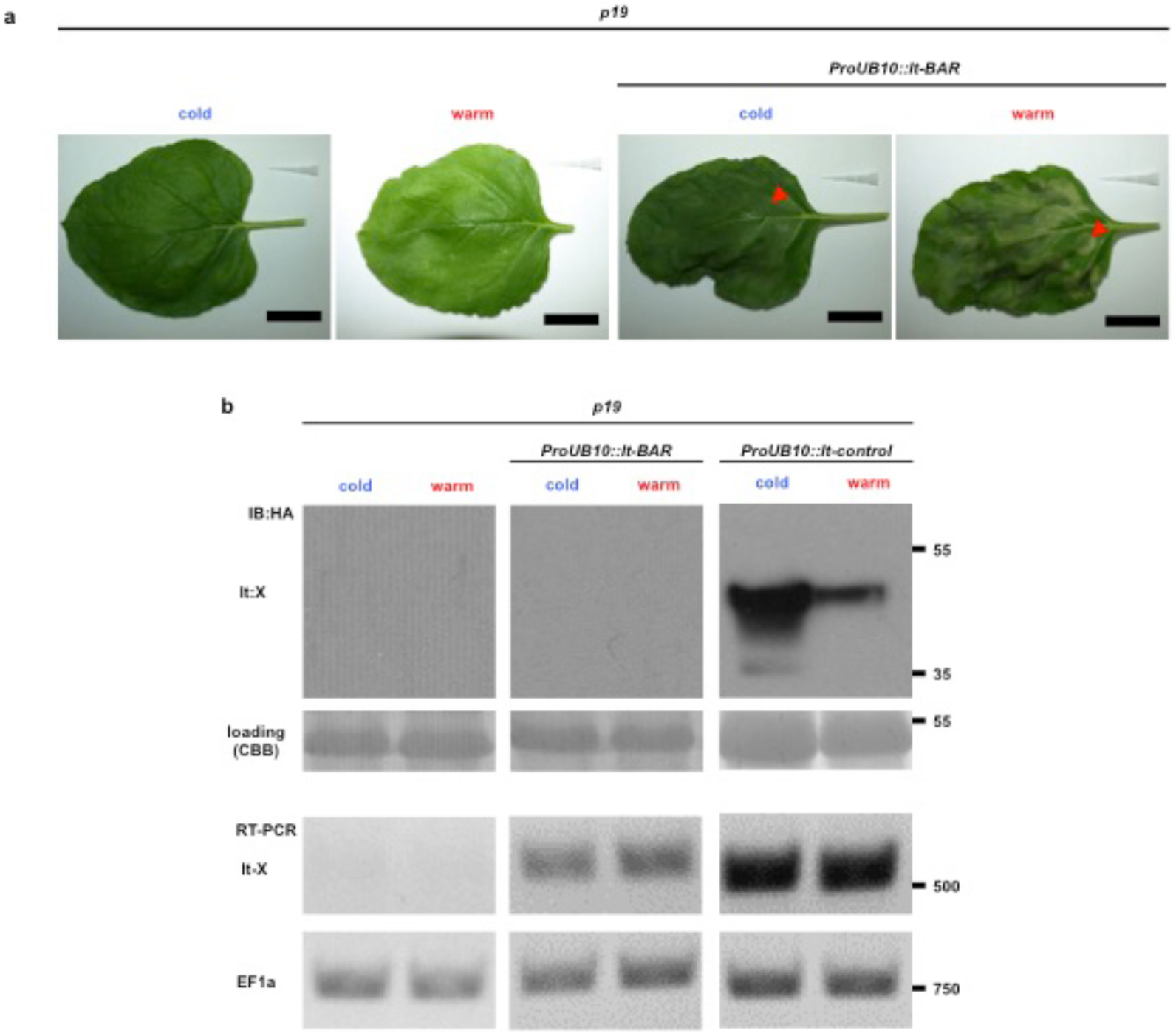
Transient expression of lt-BAR and transcript analysis. **(a)** Transient expression of lt-BAR in leaves of *N. benthamiana* leads to a temperature-independent chlorosis/necrosis phenotype. Control plants were transformed with Agrobacteria carrying the p19 plasmid only. Size bar, 3 cm. **(b)** Analysis of lt-BAR transcript and protein levels demonstrates that it is expressed but protein amounts are below the detection limit. An lt-control protein expressed from the same promoter served as a positive control during western blot analysis. Control plants were transformed with Agrobacteria carrying the p19 plasmid only.

### Conditional toxin function can be shifted and is active at ambient temperature

We then elucidated the sensitivity of the lt-BAR module. For this, plants were grown at permissive temperature, shifted after three weeks to restrictive temperatures for nine days, and finally shifted back to permissive temperature. During this process, trichomes on newly forming leaves were monitored. The trichome phenotype of shifted plants followed the temperature stimulus, exhibiting glabrous leaves at the permissive temperature, developing wild type-like leaves containing trichomes when grown under restrictive temperature, and again showing glabrous leaves when shifted back to the permissive temperature (**Figure 3a**). This shows that also the function of lt-BAR follows the temperature stimulus exhibiting efficient ON/OFF transitions as previously described for the lt-degron for non-toxic proteins (Faden et al., 2016; Dissmeyer, 2017). Next, we asked whether lt-BAR expressing plants could successfully elicit their glabrous leaf phenotype when grown at standard greenhouse conditions. Stability of lt-BAR at ambient temperature, thus the possibility to evoke a phenotype, would greatly facilitate its application as only one controlled growth environment, used for restrictive conditions, would be necessary in order to be applied successfully. Indeed, lt-BAR was able to robustly result in glabrous leaves when plants were grown in the greenhouse, suggesting sufficient stability of the fusion protein even at a temperature that is higher than the standard permissive temperature of 14°C (**Figure 3b**). Remarkably, the lt-BAR expressing plants maintain the glabrous leaf phenotype under greenhouse conditions even though the greenhouse represents a significantly less controlled environment with higher temperature fluctuations and initially higher then permissive temperatures then the previously used growth cabinets.

**Figure 3.**
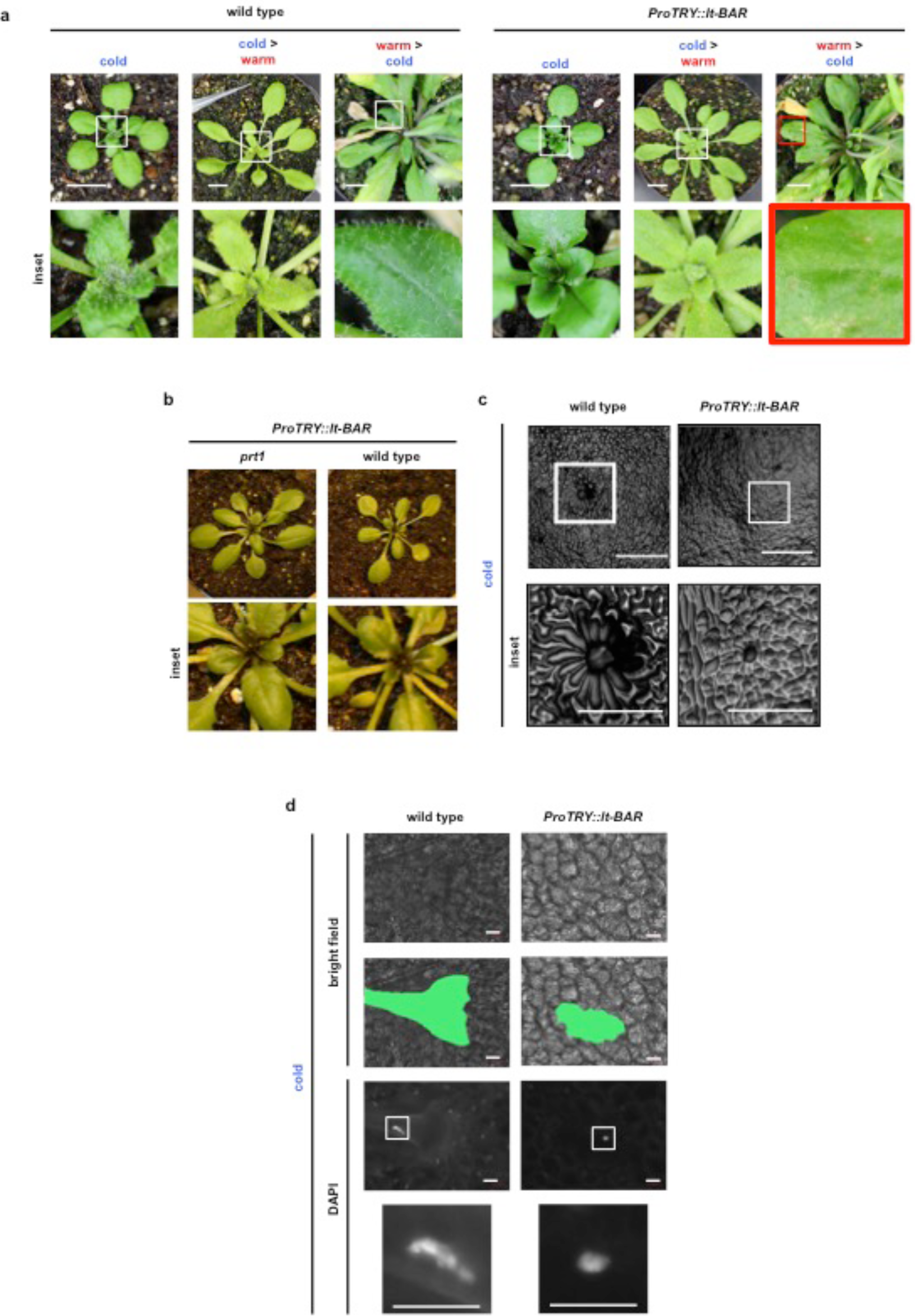
The phenotype caused by lt-BAR follows the temperature stimulus but drives cells into growth arrest rather than cell death. **(a)** Plants grown at the permissive temperature where shifted to the restrictive temperature after two weeks and shifted back to the permissive temperature after another 9 days. The glabrous leaf phenotype occurs in relation to the temperature stimulus. Size bar = 1 cm. **(b)** *ProTRY::lt-BAR* elicits a reliable “ON”-phenotype under standard greenhouse conditions in 21-day-old plants showing glabrous leaves. **(c)** Agarose imprints of the leaves of wildtype and *ProTRY::lt-BAR* plants reveal that expression of lt-BAR impairs trichome initiating cells in developing the typical single-celled trichome structure within the plant tissue. Size bar, 500 μm. **(d)** DAPI staining of leaf epidermis. The green mark in the second lane shows the surrounding of the trichome-forming cell. The unexpected shape in the wild type results from the mature trichome being pushed on the surface of the leaf by the cover slide. Size bar, 50 μm.

### Conditional toxin causes growth arrest rather than cell death

Then, we further elucidated the effect of lt-BAR on trichome development. Trichome spacing and therefore TRY activity begins in the leaf primordia (Larkin et al., 1996). Since TRY is not important for formation of the trichome itself but crucial for its spacing and patterning by suppressing trichome initiation in neighboring cells, ablation or arrest of a TRY-expressing cell could allow its expression in the neighboring cell potentially resulting in its own ablation and starting a chain reaction. To address this hypothesis, we prepared agarose imprints of the leaf surface of plants grown at the permissive temperature. Analysis revealed that the trichome forming cells went into an early arrest and were not able to form the typical trichome structure (**Figure 3c**). This could be explained through two different mechanisms. Either the *TRY* promoter is only active for a limited period which allows to express only a level of lt-BAR that is sufficient to arrest cell growth but which is not sufficient to kill the cell. Or an equilibrium is reached where synthesis of active lt-BAR and possible degradation due to a slight leakiness of the degron is reached. Such degradation at the permissive temperature has been demonstrated previously for a *lt-degron::TTG1* protein fusion (Faden et al., 2016). Also, growth arrest of a single cell, including its functional disruption, does not seem to trigger trichome formation in neighboring cells indicating that trichome spacing is a restricted process only happening for a defined period during leaf formation. To test whether the cells underwent cell death or resided in a state of growth arrest, nuclei were fluorescently stained with DAPI. All cells showed normal nuclei indicating indeed cell arrest rather than cell death (**Figure 3d**).

In very rare cases, cells were able to partially overcome the toxic effect of lt-BAR. This was indicated by started initiation and formation of a trichome in these cells (**Figure 4a**). This observation indicates that indeed the TRY promoter activity was reduced and that, potentially, cells are able to degrade the lt-BAR over an extended period of time even at the permissive temperature. However, it is noteworthy that mature trichomes where never found on any lt-BAR expressing plant at the permissive temperature. Surprisingly, the phenotype evoked by lt-BAR strikingly resembled a mutant of the transcription factor GLABRA2, the *glabra2* (*gl2*) mutant (Rerie et al., 1994; Schwab et al., 2000). The *ProTRY::lt-BAR-expressing* line shows similarly arrested cells that may grow out into basic trichome precursors (**Figures 3c,d and 4a**) as described for the *gl2* mutant. GL2 is thought to be mainly important for trichome formation, however, also playing a role in trichome patterning was discussed (Rerie et al., 1994; Hulskamp, 2004).

**Figure 4.**
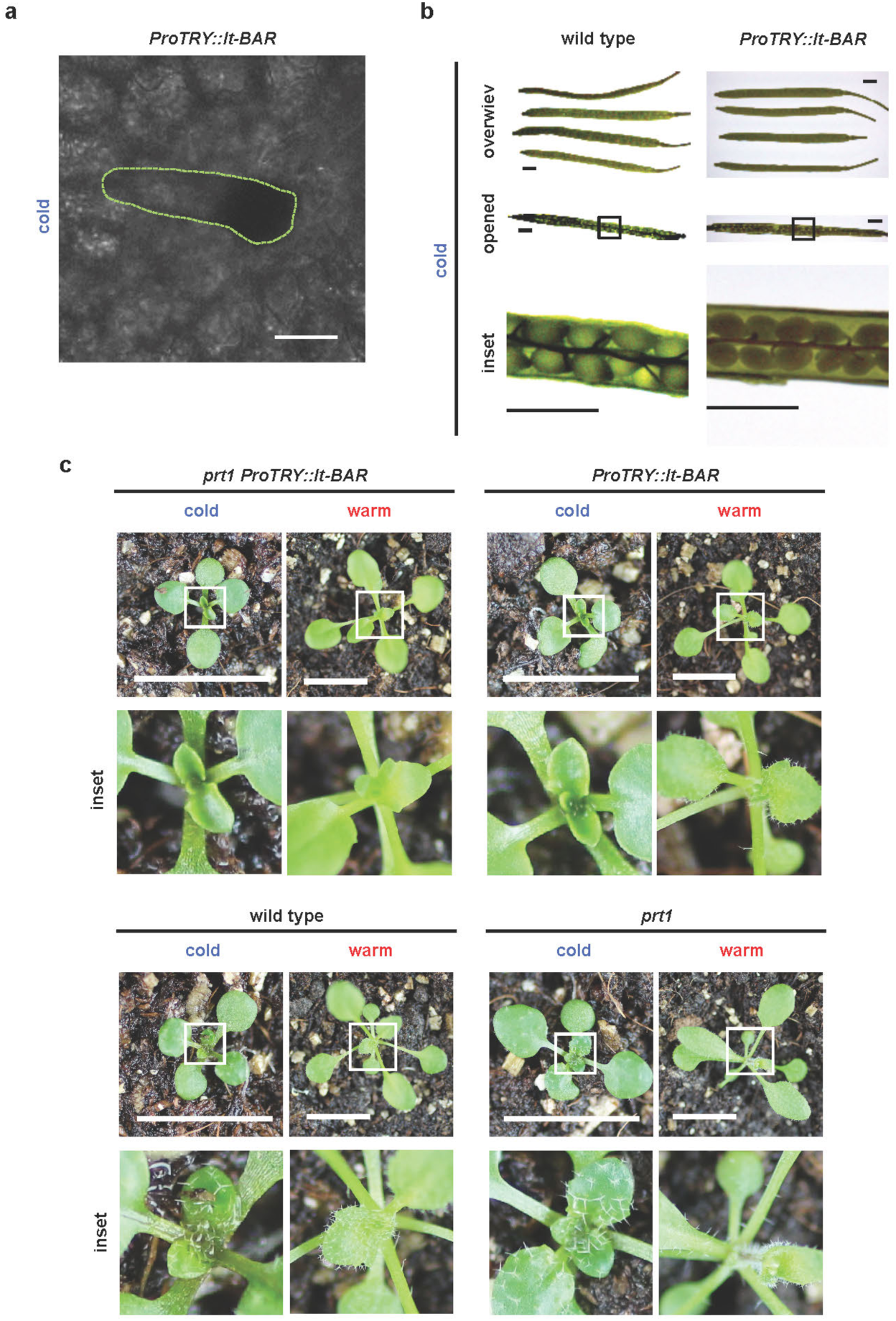
Phenotypes of lt-BAR and requirement of E3 ligase PRT1 for functionality. **(a)** Example of a trichome precursor (green-dotted shape) forming on a late leaf of a five-week-old plant expressing ProTRY::lt-BAR and grown at permissive conditions. Size bar, 50 μm. **(b)** Expression of lt-BAR in the prt1 mutant background causes temperature-independent cytotoxic effects and constitutive phenotypes of trichome ablation. Size bar, 1 cm. **(c)** Siliques of wildtype and ProTRY::lt-BAR plants. Size bar, 1 mm.

TRY is expressed in seeds (compare eFP-browser expression data; http://bar.utoronto.ca/efp/cgi-bin/efpWeb.cgi) and therefore, seed development might be affected in *ProTRY::lt-BAR* expressing plants. When plants were grown under permissive temperature, siliques appeared indistinguishable from wildtype plants (**Figure 4b**) and seeds did not show any alterations in amount, shape, and germination rate.

### N-end rule E3 ubiquitin ligase PRT1 is required for modulating lt-BAR toxicity

We recently showed that in *Arabidopsis*, lt-degron fusions, for technical reasons, are mainly degraded by the N-end rule E3 ligase PROTEOLYSIS1 (PRT1) (Faden et al., 2016) which has a high preference for aromatic N-terminal amino acids (Potuschak et al., 1998; Stary et al., 2003; Naumann et al., 2016; Dong et al., 2017; Reichman and Dissmeyer, 2017; Mot et al., 2018). To generate a stable genetic situation with constitutively stabilized lt-BAR and to test whether the observed phenotype was a result of the fusion to the lt-degron cassette or rather a response of barnase activity to the changed temperature, we introgressed *ProTRY::lt-BAR* into the *prt1* mutant. Indeed, crossing the lt-BAR into the *prt1* mutant background resulted in a complete and temperature-independent stabilization of the phenotype evoked through lt-BAR showing that presence of PRT1 and therefore altered activity of the N-end rule pathway of targeted protein degradation are responsible for the temperature-responsive phenotype (**Figure 4c**).

## DISCUSSION

Conclusively, we have shown that lt-BAR is able to efficiently mediate cell arrest and thereby influence organ fate in *Arabidopsis*. The lt-BAR is superior to many systems used due to a few of its characteristics. First of all, the control over the lt-BAR protein is easy to execute. Since it follows a temperature stimulus, stabilization as well as degradation and tuning of active protein are easy to carry out. Additionally, exogenous addition of compounds and/or other stimuli for induction of the system are not needed hence eliminating issued connected to infiltration, uneven induction of the system due to uneven perfusion of inducing agents as well as possible toxic side effects on the system by used compounds and solvents. Additionally, the temperature stimulus is easily controllable enabling for easy tuning and regulation of the system therefore rendering maintenance of the lt-BAR transgene simple and straightforward. Summing up, here we present a highly versatile and easy system for conditional organ formation.

## Acknowledgements

We thank Sylvestre Marillonnet for the clone containing the Barnase gene and fruitful discussions. This work was supported by a grant for the junior research group of the ScienceCampus Halle – Plant-based Bioeconomy to N.D., by DI 1794/3–1 and DI 1794/1 of the German Research Foundation (DFG) to N.D., by grant LSP-TP2–1 of the Research Focus Program “*Molecular Biosciences as a Motor for a Knowledge-Based Economy*” from the European Regional Development Fund (EFRE) to N.D., and Ph.D. fellowships of the Landesgraduiertenforderung Sachsen-Anhalt awarded to F.F. Financial support came from the Leibniz Association, the state of Saxony Anhalt, the DFG Graduate Training Center GRK1026 “Conformational Transitions in Macromolecular Interactions”, and the Leibniz Institute of Plant Biochemistry (IPB). This work was supported by two networks of the European Cooperation in Science and Technology (COST), namely Action BM1307 – “European network to integrate research on intracellular proteolysis pathways in health and disease (PROTEOSTASIS)” and Action FA1006 “Plant Metabolic Engineering for High Value Products (PlantEngine).”

## Author Contributions

ND initiated the project and designed the experiments; FF, SM, ND carried out the plant experiments. SM and FF performed molecular cloning; SM prepared stable plant lines and carried out transient expression tests. SM and FF did transcript analysis; FF performed protein analysis. FF and ND wrote the paper and drafted the figures.

## ORCID

Nico Dissmeyer http://orcid.org/0000–0002–4153761

